# Direction Selectivity in Naturalistic Action Observation: Distributed Representations Across the Action Observation Network

**DOI:** 10.1101/2025.10.20.683194

**Authors:** Zelal Eltaş, Murat B. Tunca, Burcu A. Urgen

## Abstract

Perceiving the direction of observed actions is critical for interpreting intentions and guiding social interaction. While direction selectivity has been extensively studied with simple stimuli such as dots, gratings, or point-light displays (PLDs), little is known about how the brain encodes direction in naturalistic, repetitive actions that are seen frequently in daily life. The present fMRI study investigated direction-selective representations during observation of complex actions performed along three bidirectional dimensions (left-right, up-down, front-back) within a 96-video stimulus set. The brain activity was analyzed using multivariate pattern analysis (MVPA) and multiple regression representational similarity analysis (RSA). MVPA revealed above-chance classification of action direction across occipital, parietal, and motor cortices, with the highest decoding in occipital, primary motor, and somatosensory regions. Crucially, RSA demonstrated that when accounting for low-level and motor features, direction information was still represented in early visual cortex, occipito-temporal areas, parietal regions, and motor-related regions. These findings indicate that action direction is represented across multiple levels of the action observation network (AON), extending from early sensory regions to higher-order parietal and frontal cortices. By using naturalistic, repetitive action videos, this study provides new evidence that the coding of action direction in the human brain is broadly distributed, reflecting the complexity of perceiving actions in everyday life. These findings suggest that direction selectivity is a core feature of the action observation network, linking basic motion processing with higher-level action understanding.

## Introduction

Perceiving and interpreting everyday actions is important in terms of understanding actions and survival of living beings (Boch et al. 2024). In humans, this ability is especially important because everyday environments are rich and complex, requiring precise discrimination of intentions, actions, objects, and motion (Lesourd et al. 2023). For example, when someone waves, one must recognize that the gesture is directed toward them, understand its social intention, and respond by waving back at the appropriate moment to engage in social interaction. Similarly, if one is chased by a dog, they must understand the intention of the dog, run as fast as they can, and be aware of surrounding objects to avoid colliding with them. This ability, especially perceiving biological motion, is both innate and universal due to its survival importance (Bardi et al. 2011; Kuhlmeier et al. 2010; Simion et al. 2008). In fact, in the literature, a specialized network in humans responsible for action perception and interpretation has been proposed consistently, called the action observation network (AON) (Buccino et al. 2001; Caspers et al. 2010; Grosbras et al. 2012). This network includes occipito-temporal regions such as the middle temporal cortex (MT+) and posterior superior temporal sulcus (pSTS), as well as parietal and frontal areas, including the anterior intraparietal sulcus (aIPS), inferior frontal cortex (IFC), and ventral premotor cortex (Caspers et al. 2010; Grosbras et al. 2012; Lesourd et al. 2023; Urgen & Saygin, 2020). These areas work hierarchically, beginning with early visual regions that extract basic visual features such as orientation and motion direction (V1/V2, MT) and extending to higher-order regions that represent action identity, meaning, and goals (Lerner et al. 2001; Neal & Kilner, 2010; Urgen et al, 2019; Urgen & Saygin, 2020; Thompson & Baccus, 2011).

Within this network, occipito-temporal regions are extensively studied and found to play a central role in more basic motion processing (Snowden & Freeman, 2004; Vaina et al. 2001). This connection is particularly important because a key feature of motion processing is direction selectivity, which has been extensively studied in early visual regions, often using macaque models and simple stimuli such as slits, single spots, or random-dot fields (Albright, 1984; Churchland et al. 2004; Hu et al. 2018; Wang & Movshon, 2015). These studies robustly revealed that there are direction-selective neurons in early visual processing regions, predominantly in MT/V5, V1, and V2 (Albright, 1984; Hu et al. 2018; Livingstone, 1998; Wang & Movshon, 2015). These neurons code motion features such as preferred direction (Albright, 1984), speed and orientation tuning (Wang & Movshon, 2015), and motion contrast sensitivity (Hu et al. 2018). Furthermore, the lateral intraparietal area (LIP) was found to encode the direction of motions according to their category (i.e., matching test and sample stimuli’s direction), while MT/V5 neurons were strongly direction selective, independent of their categories (Freedman & Assad, 2006). These results have suggested that whereas MT/V5 encodes motion direction irrespective of context, LIP transforms visual direction selectivity into more abstract representations that reflect the behavioral relevance or meaning of stimuli.

Further studies with humans using neuroimaging techniques have yielded similar results, demonstrating the importance of early visual areas on motion perception (Ajina et al. 2014; Hong et al. 2012; McKeefry et al. 1997; Prieto et al. 2007; Stoppel et al. 2011). The results of motion and direction studies revealed that V1, V2, V3, V4, and MT/V5 areas showed direction-selective responses, independent of motion types (Ajina et al. 2014; Hong et al. 2012; McKeefry et al. 1997; Prieto et al. 2007). Furthermore, in one of the studies, MT/V5 response was enhanced by increased coherence of the moving dots, and suppressed by decreased coherence of the stimuli (Stoppel et al, 2011). In sum, early visual areas, specifically MT/V5, were found to play an important role in discriminating motion direction in both macaque and human brains.

While these findings highlight the contribution of early visual regions, they rely on low-level stimuli such as dots or gratings, which provide limited insight into how the brain processes more complex, biologically relevant motion. To address this, researchers have often used point-light displays (PLDs). PLDs are created by placing dots on the joints of an actor to demonstrate motion kinematics of actions without carrying any information about actors, such as their gender, race, or appearance (Johansson, 1973). Although PLDs lack complex information regarding actions, they capture rich kinematic cues, making them useful for studying biological motion (Mather & West, 1993). A recent study has investigated the decoding of direction and velocity of biological motion, using grasping PLDs (Ziccarelli et al. 2024). PLDs have moved either to the left or to the right to perform a grasping action. The results have suggested that MT/V5, superior parieto-occipital cortex (SPOC), and medial intraparietal sulcus (mIPS) have successfully decoded the direction of grasping (Ziccarelli et al. 2024). Another studies using PLDs have shown that in the direction discrimination of a walker, the extrastriate body area (EBA) encodes the facing direction of the walker, whereas the posterior superior temporal sulcus (pSTS) and MT+ encode the walking direction of the PLDs (Vaina et al. 2001; Vangeneugden et al. 2014). Extending these findings, Grossman et al. (2010) demonstrated that the pSTS also maintains action-specific representations that are invariant to changes in size and viewpoint of PLD walkers, highlighting its role in higher-level action perception.

Nevertheless, the simplistic nature of PLDs may limit their suitability for addressing all aspects of action perception. Although PLDs are highly controlled and isolate kinematic information, they do not include the full range of cues present in daily life actions. Besides the kinematic cues, action perception in real life also involves the perception of the actors and the social cues associated with them. This is supported by previous findings that show a stronger AON activation for real-life videos compared to PLD animations (Harrison et al, 2013; Orban & Urgen, 2025). These findings show that while PLDs are valuable for isolating motor cues, they may underestimate the contribution of higher level, actor-related features present in daily life stimuli. Therefore, the crucial next step in direction selectivity studies should be to extend the findings of PLD studies to using more ecologically valid stimuli. To that end, the current study aims to address this issue by investigating direction selectivity in action perception by using real-life videos. In addition, the current study aims to build on the previous findings by using more ecologically valid actions as well. The stimuli used in early studies, both biological motion and motion direction studies, moved unidirectionally to capture the direction-specific patterns better. Yet, the real world contains much richer information and complex motion patterns. For instance, the waving action is a repeated motion to the right and left, whereas traditional direction selectivity studies use PLD actions performed towards a single direction in action studies, or use simple stimuli moving unidirectionally in motion studies. Furthermore, those repetitive bidirectional actions are very common in our daily life, such as rolling a dough (i.e., front-back), wiping the window (i.e., left-right), or shaking hands (i.e., up-down). To our knowledge, no study has investigated these types of stimuli, so whether the bidirectional actions will be represented in MT+ and pSTS as in early studies, or will be carried onto the higher levels of AON, such as the premotor cortex, remains unclear in the literature.

The current study aims to investigate direction selectivity in action perception with an fMRI study in which participants viewed eight naturalistic action videos performed in three different directions (left-right, front-back, and up-down). Unlike commonly used unidirectional moving dot stimuli in the literature, the action videos in the current study consisted of repetitive movements bidirectionally, offering more ecologically valid and complex stimulus patterns. The current study utilizes multivariate analysis techniques, including Multivariate Pattern Analysis (MVPA) and multiple regression Representational Similarity Analysis (RSA). MVPA was used to identify brain areas involved in decoding action direction. However, because MVPA does not reveal whether the decoding reflects direction information specifically or other correlated factors, RSA was applied to control for low-level and motor-related features and thereby isolate direction-specific information.

By using naturalistic videos and multivariate analysis techniques, this study aims to investigate brain regions involved in direction-selective representations of observed actions, and to generalize and extend the findings of both direction and action perception literature.

## Methods

### Participants

Twenty-seven participants (10 male, 26 right-handed; *M_age_*= 23) took part in the study. They self- reported their handedness based on their hand preference when engaging in various tasks such as writing, throwing, and using scissors. Their participation was voluntary; they did not receive any compensation. All participants reported normal or corrected-to-normal vision and had no history of neurological disorders. They filled in the fMRI compatibility and consent form prior to the study. Recruitment was conducted via social media platforms. Ethical approval for the study was obtained from the Human Research Ethics Committee of Bilkent University.

### Stimuli

Stimuli consisted of 96 action videos performed along three movement dimensions: horizontal (left-right), vertical (up-down), and depth (front-back). Each dimension included eight actions, representing two exemplars from each of four naturalistic action clusters. The four action clusters were: (1) object manipulation actions, (2) object manipulation actions with a tool, (3) self-directed actions, and (4) self-directed actions with a tool. The eight actions included shaking an object, stretching an object, painting an object with a brush, wiping an object with a cloth, rubbing oneself, scratching oneself, brushing oneself with a facial cleansing brush, and drying oneself with a cloth (Table 1). Actions were selected based on their ability to be performed repetitively and naturally in each movement dimension to ensure ecological validity. To increase stimulus variability, two versions of each action were created by altering either the object, the body part involved, or the tool used. All actions were performed by both male and female actors.

**Table 1.** A table showing the action types, action clusters, whether a tool is used while performing the actions, and the two variations of the actions in the study.

| Action Name | Action Cluster | Tool Used | Variations |
| --- | --- | --- | --- |
| Shaking | Object manipulation | None | Water bottle, cola bottle |
| Stretching |  |  | Yellow resistance band, pink resistance band |
| Painting | Object manipulation with a tool | Painting brush | Purple acrylic paint, green acrylic paint |
| Wiping |  | Cloth | Red mug, cardboard box |
| Rubbing |  |  | Forehead area, neck area |
| Scratching | Self-directed | None | Forehead area, neck area |
| Drying | Self-directed with a tool | Cloth | Mouth area, forehead area |
| Brushing |  | Facial cleansing brush | Cheek area, forehead area |

Objects and tools used in the stimuli included plastic bottles, resistance bands, a cardboard box, a mug, acrylic paints, a brush, and a cloth (Table 1). For self-directed actions, actors either used their bare hands or applied tools such as a facial cleansing brush or cloth.

All videos were filmed against a gray wall and a light yellow table. Only the actor’s upper body was visible. Actors maintained a downward gaze toward the table and used their right hand for all actions. Actions were performed while facing the camera directly, with the torso and head aligned with the lens. During filming, object/tool/body part positions were adjusted to suit each movement dimension without changing the torso and face position. For instance, while the front part of the forehead is scratched for the up-down and left-right action, the side of the forehead is used for back-front to represent the depth information clearly.

Each action, with both of its variations, was performed by both male and female actors in all three directions. This resulted in 96 video stimuli in total (8 actions × 2 variations × 2 actors × 3 directions), each lasting 3 seconds.

### Experimental Design

#### Procedure

The experiment was programmed using MATLAB (The MathWorks Inc., 2022) and the Psychtoolbox extension (Brainard, 1997; Pelli, 1997; Kleiner et al, 2007). Participants were informed about the study, experimental setup, and signed consent forms before participating. Prior to the functional scans, a high-resolution 6-minute anatomical scan was applied, followed by the experiment.

The experiment consisted of eight runs, each 8 minutes long and comprising 48 trials. Each trial presented a unique combination of action, direction, and actor. Odd-numbered runs (i.e., 1, 3, 5, 7) featured the first variation of each action; even-numbered runs (i.e., 2, 4, 6, 8) featured the second variation. Thus, the full set of 96 videos was divided evenly across the eight runs, in a way that every combination of action, direction, and actor was seen in a run. Each movement direction was shown 16 times per run, totaling 128 repetitions per participant across the experiment. The full session lasted approximately 70 minutes, including 6 minutes of anatomical scans.

Each run began with a 10-second black fixation cross (i.e., rest), followed by trials (Fig. 1). Each trial started with a black fixation cross displayed for a randomly determined duration between 3 and 5 seconds, which was subsequently followed by a 3-second action video (800 x 450 pixels, 25 FPS). During the video, the fixation cross turned into a white dot for visibility, as actors wore black clothing. To minimize the eye movement of the participants, videos were slightly shifted (i.e., a minimum of 10, a maximum of 150 pixels) on the screen to align the relevant action area (e.g., forehead or object) with the fixation point, consistent with previous work (Urgen et al, 2021). For instance, for the rubbing action, a video is slightly shifted downwards to align the fixation point with the hand position of the actor, while for the painting action, it is shifted upwards towards the box.

**Fig. 1.**
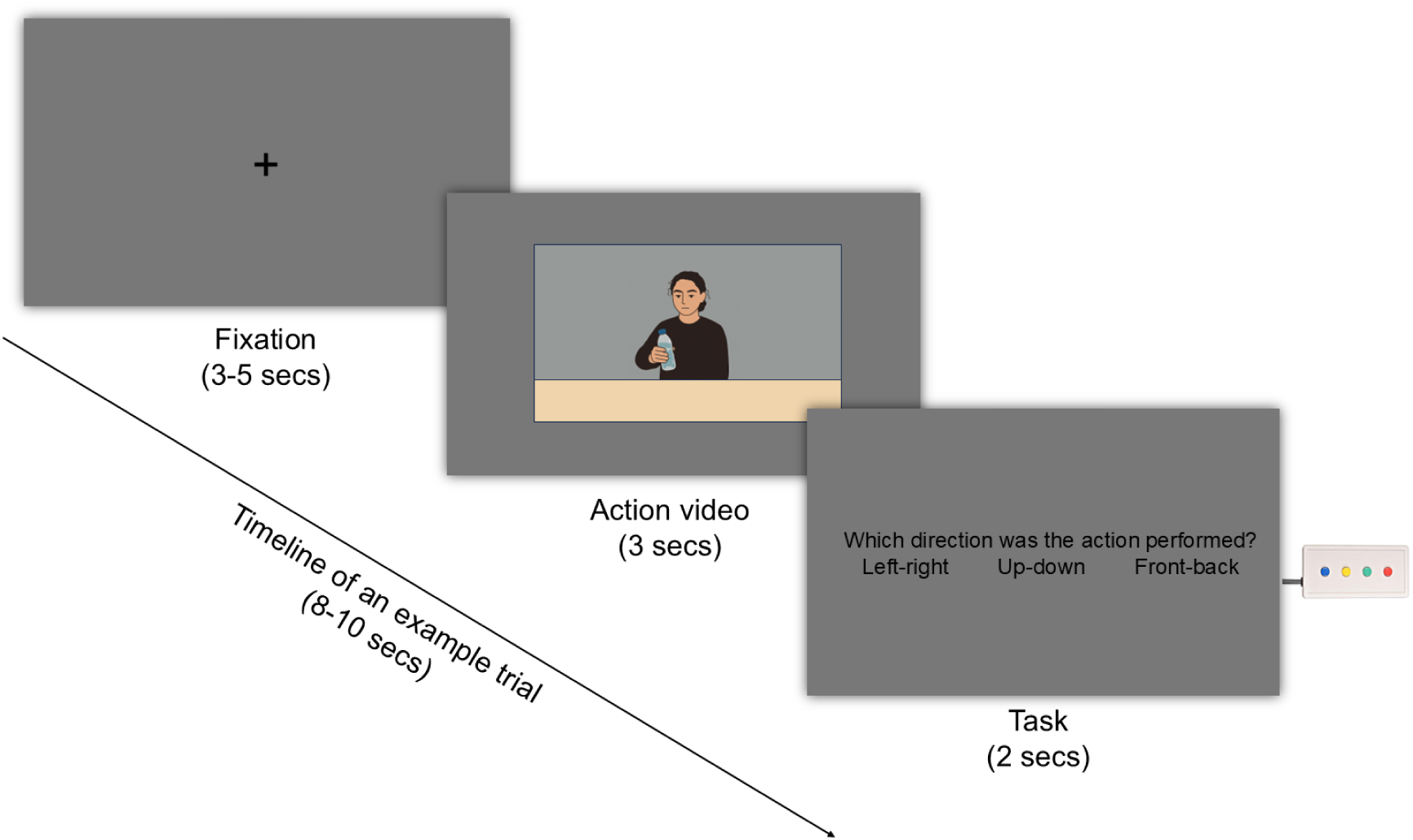
Example timeline of one trial in the fMRI experiment. Trials began with a fixation cross, followed by an action video and a task requiring participants to indicate the direction of the performed action. Participants gave responses by pressing the buttons on the button box, shown on the right side of the figure. The action video is represented as a drawing for illustrative purposes; the actual study used real action videos.

After the video, a task screen appeared for 2 seconds. Participants indicated the direction of the movement using a button box with three options: left–right (blue button), up–down (yellow button), or back–front (green button) (Fig. 1). The red button in the button box has not been used. Responses to text appearing on the left side of the screen (i.e., left-right) were made with the left thumb (blue button), whereas responses to text appearing on the right side of the screen (i.e., front-back) were made with the right thumb (green button). Participants were instructed to press the middle (yellow) button with their right thumb. However, inspection of contralateral somatosensory activation in the GLM analysis (see Supplementary File 1- Figure S1 for detailed explanation) indicated that not all participants followed this instruction consistently: 16 participants predominantly used their left hand, whereas 9 participants predominantly used their right hand to press the middle button. After the response, no feedback was provided. The response screen remained visible for the full 2 seconds regardless of response time. Each run concluded with another 10-second rest period.

The purpose of task was to ensure sustained alertness throughout the fMRI session and to confirm that participants attended to the directions of the actions rather than unrelated visual features of the videos. Participants demonstrated high task engagement, with a mean accuracy of 97%.

#### fMRI Data Acquisition

Data were acquired at the National Magnetic Resonance Research Center (UMRAM) at Bilkent University using a 3T Siemens TimTrio MRI scanner with a 32-channel head coil. Foam cushions were placed around the head, neck, and under the legs to minimize motion. Stimuli were presented on an MR-compatible 32-inch LED monitor (Samsung, 60 Hz refresh rate, 1920×1080 resolution) located 168 cm from the participant’s view. A mirror mounted on the head coil allowed participants to view the screen. Prior to functional scanning, a high-resolution T1-weighted anatomical image was collected (TE = 2.92 ms; TR = 2.6 s; flip angle = 12°; acceleration factor = 2; resolution = 1 mm³; 176 sagittal slices). Functional images were acquired using EPI sequences (TE = 30 ms; TR = 2 s; flip angle = 90°; FOV = 240 mm; matrix = 96×96; 49 slices; 2.5 mm slice thickness). Each run began with three dummy scans to allow for magnet stabilization.

### fMRI Data Analysis

#### Preprocessing

All fMRI data were preprocessed using *fMRIPrep* (version 20.2.6; Esteban et al., 2018), a standardized and widely used preprocessing pipeline based on *Nipype* (version 1.7.0; Gorgolewski et al., 2022).

##### Anatomical preprocessing

Anatomical images consisted of two T1-weighted scans per participant. These images were corrected for intensity non-uniformity using N4BiasFieldCorrection (ANTs 2.3.3) and skull-stripped using an ANTs-based workflow with the OASIS30ANTs template. Brain tissue segmentation into gray matter (GM), white matter (WM), and cerebrospinal fluid (CSF) was performed using FSL FAST. A T1-weighted reference image was created by registering the two corrected T1-weighted images, and cortical surface reconstruction was carried out using FreeSurfer (version 6.0.1). Spatial normalization to MNI space (MNI152NLin2009cAsym) was performed using nonlinear registration with ANTs.

##### Functional preprocessing

For each of the eight functional runs per participant, a reference BOLD volume was generated and skull-stripped using *fMRIPrep*’s internal workflow. Functional images were co-registered to the T1-weighted reference using boundary-based registration implemented in FreeSurfer with six degrees of freedom. Head motion was estimated using FSL MCFLIRT, and slice-timing correction was performed using AFNI 3dTshift. Functional data were resampled both in native space and in standard MNI space.

Several confound regressors were computed, including framewise displacement (FD), DVARS, and global signals extracted from CSF, WM, and whole-brain masks. Component-based noise correction (CompCor) was performed by extracting principal components from high-variance voxels (tCompCor) and from anatomical CSF and WM masks (aCompCor). For each CompCor decomposition, components explaining up to 50% of variance were retained. Motion parameters, their temporal derivatives, and quadratic terms were included as additional confounds. Volumes exceeding thresholds of 0.5 mm FD or 1.5 standardized DVARS were flagged as motion outliers.

All spatial resampling was performed in a single interpolation step using ANTs with Lanczos interpolation to minimize smoothing. Surface-based resampling was performed using FreeSurfer tools. Additional internal operations relied on *Nilearn* (version 0.6.2; RRID:SCR_001362).

##### Exclusions

Runs with excessive head motions were excluded from the analysis based on the head motion calculations of fMRIPrep. The excessive head motions represented translations more than 1.5 mm or rotations more than 1.5°. One participant was excluded from the analysis because he had either excessive rotation or translation head motions in all runs. Another participant was excluded because of a code malfunction during the fMRI scan. A total of two participants (2 male, 1 left-handed, *M_age_* = 23) were excluded from the analysis.

From the remaining twenty-five participants (8M, all right-handed *M_age_* = 23), a total of 11 runs were excluded because of excessive head motions: two runs each from three participants and one run each from five participants. One participant’s data were collected in two sessions (four runs in each) due to technical problems with the fMRI machine.

### Univariate Analyses

Univariate analyses were conducted using the General Linear Model (GLM) implemented in Statistical Parametric Mapping 12 (SPM12; Ashburner et al. 2021) in MATLAB (The MathWorks Inc., 2022). Two separate GLMs were specified to support multivariate analyses. The first GLM was constructed for multivariate pattern analysis (MVPA) and included 12 regressors: 3 action directions (right–left, up–down, front–back), inter-stimulus intervals, response periods, rest periods, and 6 motion parameters (3 translations and 3 rotations). Beta maps of action direction periods were used in decoding analyses.

The second GLM was designed for representational similarity analysis (RSA) and included 105 regressors: 96 action videos, inter-stimulus intervals, response periods, rest periods, and 6 motion parameters (3 translations and 3 rotations). The resulting beta maps for each action video period were used as input for searchlight-based RSA.

### Searchlight Multivariate Pattern Analysis (MVPA)

MVPA applies machine learning to fMRI data by training classifiers to detect reproducible spatial activity patterns that distinguish experimental conditions (Mahmoudi et al. 2012). While GLM identifies average activation differences, MVPA allows for the localization of brain regions discriminating between movement directions more sensitively and robustly. The analysis was performed using The Decoding Toolbox (TDT; Hebart et al. 2015) in MATLAB (The MathWorks Inc., 2022). Preprocessed beta images from GLM analysis were used as input. A whole-brain searchlight analysis with a sphere radius of 4 mm was conducted using a linear support vector machine (SVM) classifier and leave-one-run-out cross-validation. Classification was performed across three action directions (right-left, up-down, front-back), and model performance was quantified using accuracy-minus-chance and confusion matrices.

Group-level statistical inference was conducted using threshold-free cluster enhancement (TFCE) with permutation testing, corrected for multiple comparisons, as implemented in CoSMoMVPA (Smith & Nichols, 2009; Oosterhof et al. 2016). Accuracy-minus-chance maps from each subject were entered into a group-level analysis. A TFCE-based Monte Carlo simulation with 10,000 permutations was performed to evaluate the spatial consistency of decoding effects across participants. Z-statistic maps were generated with a correction for multiple comparisons, and a one-tailed threshold of *p* < 0.05 (*z* = 1.65) was applied. The group mean of the accuracy-minus-chance maps was calculated and thresholded with the TFCE map created to obtain significant clusters.

The brain maps were visualized with the BrainNet viewer (Xia et al. 2013). The resulting brain MNI coordinates were labeled by using Automated Anatomical Labelling Atlas 3 (AAL3; Rolls et al. 2020), implemented in MRIcroGL software (Rorden, 2025).

### Searchlight Model-based Representational Similarity Analysis

Six model RDMs (Representational Dissimilarity Matrices) were created to identify the hypothesized dissimilarities between action videos. The RDMs were 96×96 in size, representing the pairwise dissimilarities of 96 action videos. The RDMs included four categorical models (direction, action type, motor response, and tool), one conceptual model (Hierarchical Model and X -HMAX-C1), and one random model (Fig. 2). The motor response model was defined in a subject-specific manner based on the thumb used to press the middle button on the button box. For participants who responded with their right thumb, the right-thumb motor response RDM was included in the regression; for participants who responded with their left thumb, the corresponding left-thumb motor response RDM was used instead. In both cases, the remaining non-motor model RDMs were identical, resulting in the same total number of predictors (six) for all participants.

**Fig. 2.**
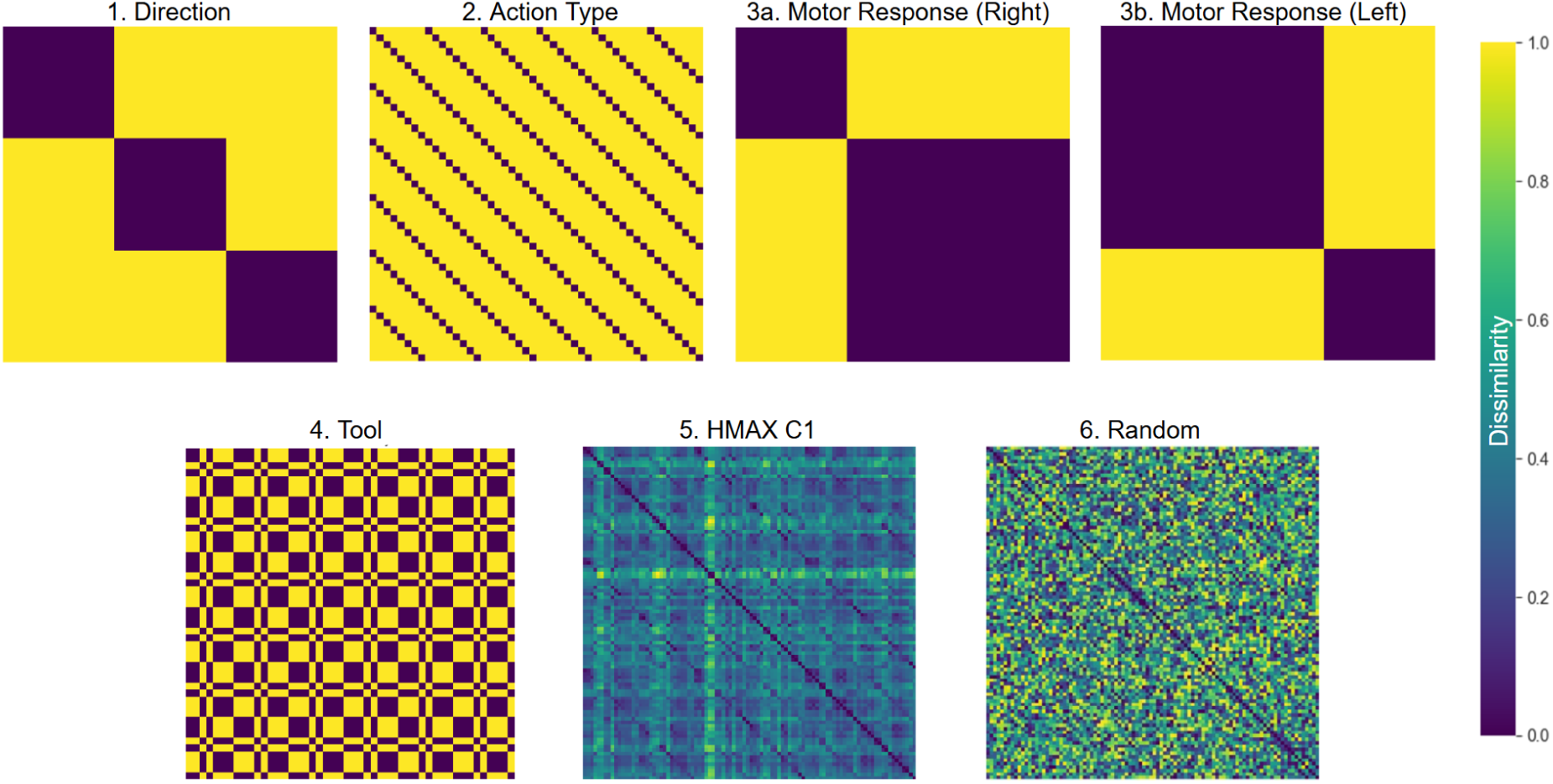
Six model RDMs used in the RSA: direction, action type, motor response (changes subject-specifically according to the thumb for middle button press), tool, HMAX C1 and random models. The matrices are 96×96, representing each stimulus pair. 0 indicates that the two stimuli are the same, while 1 indicates that the two stimuli are very different.

The direction model represents the direction in which the action is performed (Figure 3.1). The action type model represents the actions used in the videos (Figure 3.2). In the motor response model, the thumb used by participants to respond was coded to eliminate activities related to potential motor preparation. For participants who pressed the middle button with their right thumb, motor response (right) model was used (Figure 3.3a). For participants who pressed the middle button with their left thumb, motor response (left) model was used (Figure 3.3b). The tool model represents whether a tool was involved in the action (i.e., cloth, facial cleansing brush, or paint brush) (Figure 3.4). In all six models, a value of 0 indicates that two stimuli belong to the same category, and a value of 1 indicates that they belong to different categories. The fifth and more complex HMAX C1 model is an early-level component of a biologically-inspired visual processing model (Bileschi et al. 2012; Riesenhuber & Poggio, 1999; Serre et al, 2007). It models the low/mid-level features of stimuli (such as edge, shape, color, motion) by imitating complex cell responses at the V1–V2 level in the brain. Data for each stimulus was obtained from this model, the correlation between the data for every two stimuli was calculated, and this was encoded into the dissimilarity matrix using the (1 - correlation) method (Figure 3.5). A value of 0 indicates that the low/mid-level visual features of the two stimuli are the same, while a value of 1 indicates they are completely different. Finally, the random model was created by generating a random 96×96 matrix in MATLAB, with its diagonal values set to 0 and its second half being the mirror image of the first half (Figure 3.6). To assess multicollinearity among the representational dissimilarity matrices (RDMs), the Variance Inflation Factor (VIF) for each model was calculated. Results indicated low risk of multicollinearity for all models (Direction: 1.67, Action: 1.24, Motor: 1.67, HMAX: 1.08, Tool: 1.17, Random: 1.01). These representational dissimilarity matrices (RDMs) were included as input in a multiple regression analysis to determine the unique contribution of each variable in explaining neural patterns, while controlling for the effects of other predictors. Multiple regression representational similarity analysis (RSA) was performed using CoSMoMVPA (Oosterhof et al. 2016). For each subject, beta estimates corresponding to individual action videos obtained from first-level GLM analyses were used to represent condition-specific neural activation patterns. Voxels with zero signal across all samples and any additional non-contributing data were removed for clarity. To control for run-wise variability, data were z-scored within each run. Afterwards, activation patterns for each condition were averaged across runs to achieve a single, stable multivoxel pattern per condition. The resulting dataset was used as input for the representational similarity analysis.

**Fig. 3.**
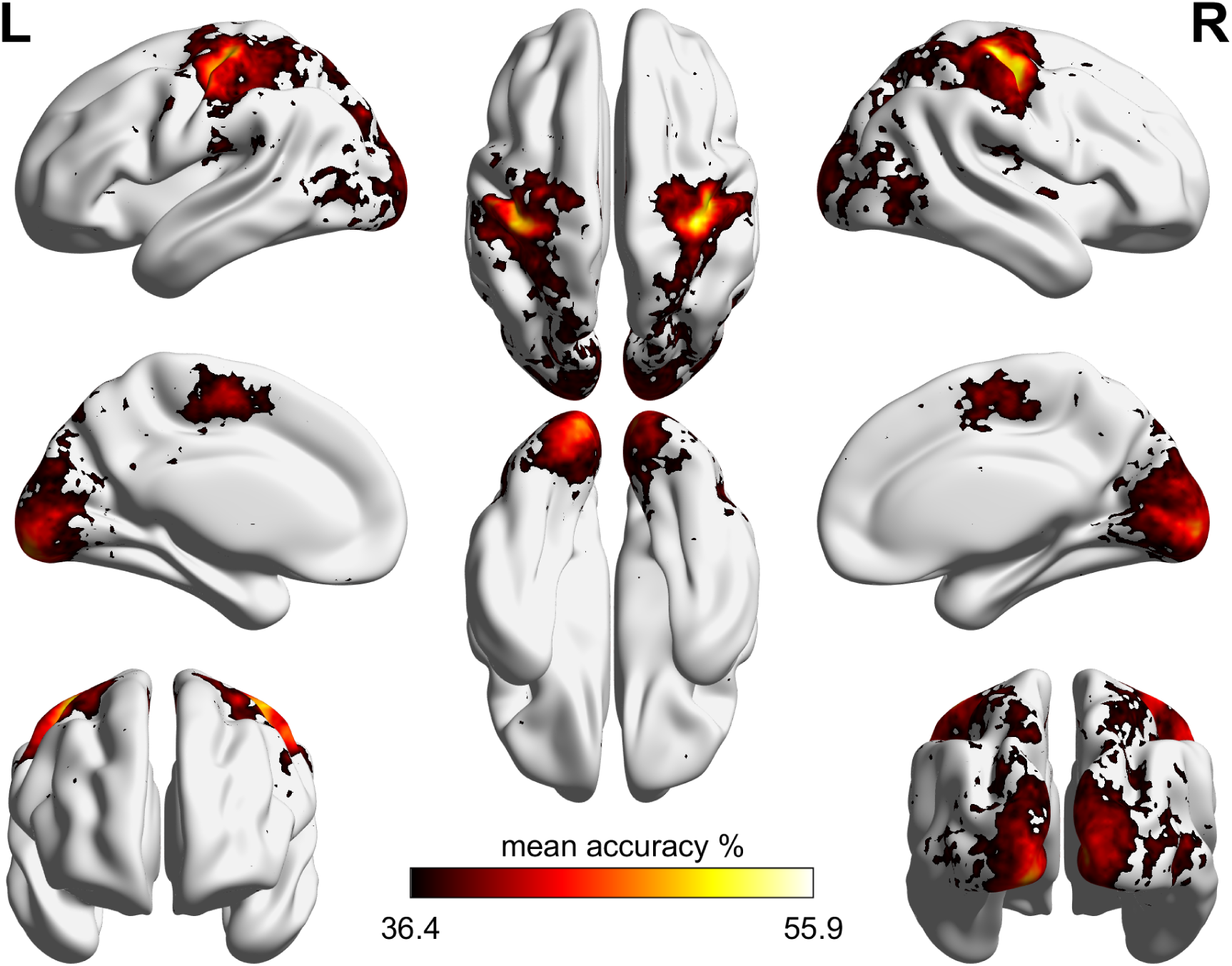
Group-level mean MVPA decoding accuracy results, thresholded with TFCE-based Monte Carlo simulation with 10,000 permutations (*p* < 0.05, *z* = 1.65). Colorbar shows mean decoding accuracy percentages (chance level = 33.3%).

For RSA, a spherical searchlight with a radius of 4 mm was applied across the whole brain to extract local multivoxel patterns. Within each searchlight, a neural representational dissimilarity matrix (RDM) was computed using squared Euclidean distance between activation patterns. This neural RDM was then entered into a multiple regression model, where it was predicted by the aforementioned six model RDMs. The regression was performed locally at each searchlight location, yielding beta maps that reflected the unique contribution of each model in explaining local neural representational structure while accounting for the influence of other predictors.

Group-level statistical inference was conducted using threshold-free cluster enhancement (TFCE) with permutation testing, corrected for multiple comparisons, as implemented in CoSMoMVPA (Oosterhof et al. 2016). For each model, subject-level beta maps from the multiple regression RSA were entered into a one-sample test against zero. A TFCE-based Monte Carlo simulation with 10,000 permutations was performed to assess the spatial consistency of RSA effects across participants. Z-statistic maps were generated for each model, using a one-tailed threshold of *p* < 0.05 (*z* = 1.65). The t-test results were thresholded with the TFCE maps to obtain the significant clusters.

The brain maps were visualized with the BrainNet viewer (Xia et al. 2013). The resulting brain MNI coordinates were labeled by using Automated Anatomical Labelling Atlas 3 (AAL3; Rolls et al. 2020) implemented in MRIcroGL software (Rorden, 2025).

## Results

### Searchlight Multivariate Pattern Analysis (MVPA)

For the action direction, a three-way searchlight MVPA analysis with a 4 mm spherical radius was performed (Fig. 3). Group-level mean accuracy maps were computed across participants and thresholded with significant clusters obtained from threshold-free cluster enhancement, corrected for multiple comparisons (TFCE; *p* < .05, 10,000 permutations, thresholded at *z* = 1.65). All thresholded areas showed above-chance decoding accuracy (chance level = 33.3%). Subject-based accuracy-minus-chance maps can be accessed from Supplementary File 2.

The MVPA analysis revealed significant decoding accuracies across a distributed set of occipital, parietal, and motor-related regions (Fig. 3). In the early visual cortex, strong effects were observed bilaterally in the calcarine sulcus (R: 47.6%; L: 45.6%, *p* < .001) and the lingual gyrus (R: 41.1%; L: 43.3%, *p* < .001). Additional occipito-temporal contributions included the superior occipital gyrus (SOG; R: 39.6%; L: 38.8%, *p* < .001), the inferior occipital gyrus (L IOG: 48.8%, *p* < .001), bilateral middle occipital gyrus (MOG; R: 39.4%; L: 41.2%, *p* < .001), the right lateral occipital cortex (R LOC: 39.4%, *p* < .001), and bilateral MT/V5 (R MT: 41.6%; L MT: 38.3%, *p* < .001).

Parietal regions also showed significant decoding, including the left supramarginal gyrus (40.1%, *p* < .001), bilateral superior parietal lobule (SPL; R: 41.2%; L: 43.3%, *p* < .001), and the left inferior parietal lobule (L IPL: 40.6%, *p* < .001).

In motor-related cortices, robust decoding accuracies were found bilaterally in the precentral gyrus (primary motor cortex, M1; R: 54.8%; L: 50.8%, *p* < .001) and postcentral gyrus (primary somatosensory cortex, S1; R: 48.3%; L: 49.5%, *p* < .001). Significant contributions were also observed in the supplementary motor area (SMA; R: 42.1%; L: 44.1%, *p* < .001).

Overall, these findings demonstrate that direction decoding can be reliably achieved across early visual, occipito-temporal, parietal, and motor cortices, with the highest accuracies observed in sensorimotor regions, particularly the precentral and postcentral gyri. Table 2 presents the peak coordinates of the resulting clusters together with their decoding accuracy percentages and *p*-values thresholded with the TFCE method, corrected for multiple comparisons (*p* < .05, *z* = 1.65).

**Table 2.** Decoding results of MVPA with peak coordinates, region labels, decoding accuracy percentage, and *p*-values. R: right, L: left, SOG: superior occipital cortex, IOG: inferior occipital gyrus, MOG: middle occipital gyrus, LOC: lateral occipital cortex, MT: middle temporal area (V5), SPL: superior parietal lobule, IPL: inferior parietal lobule, S1: primary somatosensory cortex, M1: primary motor cortex, SMA: supplementary motor area. The labels of the regions were constructed by using the AAL3 atlas.

|  | MNI Coordinates |  |  |  |  |
| --- | --- | --- | --- | --- | --- |
| Region (AAL3) | x | y | z | decoding accuracy % | $p$ |
| R Calcarine | 10 | -94 | -2 | 47,6 | <.001 |
| L Calcarine | -10 | -94 | -5 | 45,6 | <.001 |
| R Lingual | 13 | -77 | -5 | 41,1 | <.001 |
| L Lingual | -17 | -82 | -12 | 43,3 | <.001 |
| R SOG | 21 | -91 | 29 | 39,6 | <.001 |
| L SOG | -17 | -89 | 40 | 38,8 | <.001 |
| L IOG | -18 | -92 | -10 | 48,8 | <.001 |
| R MOG | 45 | -81 | 6 | 39,4 | <.001 |
| L MOG | -48 | -72 | 0 | 41,2 | <.001 |
| R LOC | 51 | -69 | -8 | 39,4 | <.001 |
| R MT/V5 | 48 | -58 | 6 | 41,6 | <.001 |
| L MT/V5 | -47 | -67 | 11 | 38,3 | <.001 |
| L Supramarginal | -53 | -42 | 25 | 40,1 | <.001 |
| R SPL | 41 | -44 | 62 | 41,2 | <.001 |
| L SPL | -37 | -44 | 65 | 43,3 | <.001 |
| L IPL | -42 | -45 | 56 | 40,6 | <.001 |
| R Postcentral (S1) | 38 | -27 | 62 | 48,3 | <.001 |
| L Postcentral (S2) | -39 | -27 | 59 | 49,5 | <.001 |
| R Precentral (M1) | 45 | -16 | 62 | 54,8 | <.001 |
| L Precentral (M1) | -40 | -24 | 63 | 50,8 | <.001 |
| R SMA | 2 | -13 | 60 | 42,1 | <.001 |
| L SMA | -1 | 14 | 60 | 44,1 | <.001 |

### Model-based Representational Similarity Analysis

To determine the direction-specific areas by accounting for low-level or motor features, multiple regression RSA with a searchlight radius of 4 mm was conducted. For group-level analysis, a one-sample t-test across the participants was computed and then thresholded with significant clusters obtained from threshold-free cluster enhancement, corrected for multiple comparisons (TFCE; *p* < .05, 10,000 permutations, thresholded at *z* = 1.65). For anatomical labeling, along with AAL3 atlas, a meta-analysis results related to action observation was also used to ensure more specific labeling (Caspers et al. 2010).

When accounting for low-level visual (HMAX C1), motor, action, and random models, the direction model revealed significant representational similarity across a broad network of regions (Fig. 4a).

**Fig. 4.**
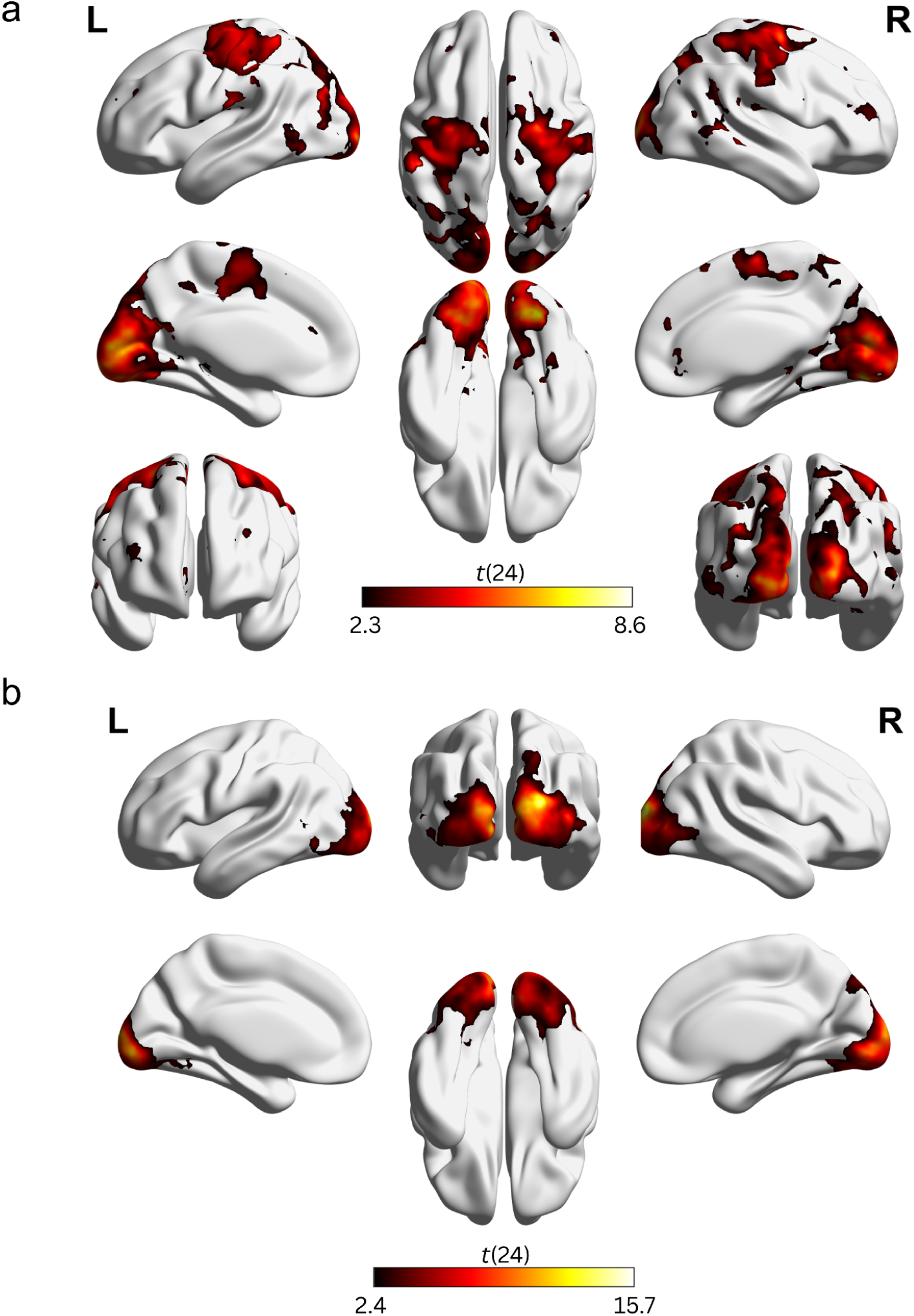
One-sample t-test multiple regression RSA results, thresholded with TFCE-based Monte Carlo simulation with 10,000 permutations (*p* < 0.05, *z* = 1.65). Colorbar shows t-values. a) Result of the direction model when accounting for HMAX C1, motor, action, and random models. b) Result of the HMAX C1 model when accounting for direction, action, motor, and random models.

In early visual regions, bilateral Calcarine (R: *t*(24) = 8.5; L: *t*(24) = 7.7, *p* < .001), left Cuneus (L: *t*(24) = 5.9, *p* = .001) and the right Lingual Gyrus (R: *t*(24) = 5.5, *p* < .001) exhibited significant contributions. Additional representation was detected throughout the occipital and temporal cortex in bilateral Fusiform Gyrus (R: *t*(24) = 7.5; L: *t*(24) = 7.1, *p* < .001), the left Middle Occipital Gyrus (MOG; L: *t*(24) = 7.9, *p* < .001), left Lateral Occipital Cortex (LOC; L: *t*(24) = 4.5, *p* < .001), bilateral Superior Occipital Gyrus (SOG; R: *t*(24) = 4.7; L: *t*(24) = 5.4, *p* < .001), the left Inferior Occipital Gyrus (IOG; L: *t*(24) = 7.0, *p* < .001), the left Inferior Temporal Gyrus (ITG; L: *t*(24) = 3.6, *p* < .001), and bilateral MT (R: *t*(24) = 4.1, *p* = .006; L: *t*(24) = 3.1, *p* < .001).

Within the parietal cortex, information was represented in the left Inferior Parietal Sulcus (IPS; L: *t*(24) = 6.8, *p* < .001), bilateral Superior Parietal Lobule (SPL; R: *t*(24) = 4.0, *p* = .005; L: *t*(24) = 5.0, *p* = .006), and bilateral Supramarginal Gyrus (R: *t*(24) = 5.8, *p* < .001; L: *t*(24) = 5.2, *p* = .008).

Motor and somatosensory areas also showed considerable involvement, including bilateral Precentral Gyrus (M1; R: *t*(24) = 4.8; L: *t*(24) = 4.9, *p* < .001), bilateral Postcentral Gyrus (S1; R: *t*(24) = 3.7; L: *t*(24) = 4.4, *p* < .001), and bilateral Supplementary Motor Area (SMA; R: *t*(24) = 5.1; L: *t*(24) = 3.9, *p* < .001). Further motor contributions were observed in the bilateral dorsal Premotor Cortex (dPMC; R; *t*(24) = 6.4, *p* < .001; L; *t*(24) = 3.7, *p* < .001).

Regions within the frontal cortex and related structures included bilateral Middle Frontal Gyrus (MFG; R: *t*(24) = 3.4, *p =* .046; L: *t*(24) = 3.9, *p =* .028).

Overall, these results indicate that direction information is represented across a broad cortical network, spanning early visual areas, higher-order occipito-temporal regions, parietal cortex, and motor-related regions, suggesting a distributed representational basis for action direction in the human brain. Table 3 shows the peak MNI coordinates of these regions, their anatomical labeling using the AAL3 atlas, *t*- and *p*-values.

**Table 3.** The region labels, peak MNI coordinates, *t*- and *p*-values of the results of multiple regression RSA results of the direction model. R: right hemisphere, L: left hemisphere, MOG: middle occipital cortex, LOC: lateral occipital cortex, SOG: superior occipital cortex, MT: middle temporal area, ITG: inferior temporal gyrus, IPS: inferior parietal sulcus, SPL: superior parietal lobule, S1: primary somatosensory cortex, M1: primary motor cortex, SMA: supplementary motor area, dPMC: dorsal premotor area (Brodmann area 6), MFG: middle frontal gyrus.

| Region (AAL3) | MNI Coordinates | | | $t(24)$ | $p$ |
| --- | --- | --- | --- | --- | --- |
|  | x | y | z |  |  |
| R Calcarine | 18 | -93 | 2 | 8.5 | <.001 |
| L Calcarine | -18 | -86 | 0 | 7.7 | <.001 |
| R Lingual | 15 | -57 | -9 | 5.4 | <.001 |
| L Cuneus | -16 | -62 | 23 | 5.6 | .001 |
| R Fusiform | 25 | -80 | -11 | 7.5 | <.001 |
| L Fusiform | -26 | -76 | -6 | 7.1 | <.001 |
| R Supramarginal | 55 | -46 | 27 | 5.8 | <.001 |
| L Supramarginal | -58 | -21 | 20 | 5.2 | .008 |
| L MOG | -20 | -94 | -2 | 7.9 | <.001 |
| L LOC | -39 | -75 | 23 | 4.5 | <.001 |
| R SOG | 23 | -62 | 35 | 4.7 | <.001 |
| L SOG | -20 | -72 | 23 | 5.4 | <.001 |
| L IOG | -20 | -94 | -6 | 7 | <.001 |
| R MT | 55 | -49 | 2 | 4.1 | .006 |
| L MT | -50 | -52 | 5 | 3.4 | <.001 |
| L ITG | -45 | -63 | -4 | 3.6 | <.001 |
| L IPS | -56 | -32 | 48 | 6.8 | <.001 |
| R SPL | 39 | -44 | 63 | 4 | .005 |
| L SPL | -33 | -50 | 69 | 5 | .006 |
| R S1 | 29 | -42 | 58 | 3.7 | <.001 |
| L S1 | -40 | -29 | 59 | 4.4 | <.001 |
| R M1 | 29 | -18 | 67 | 4.8 | <.001 |
| L M1 | 42 | -30 | 53 | 4.9 | <.001 |
| R SMA | 4 | -10 | 61 | 5.1 | <.001 |
| L SMA | -8 | -9 | 56 | 3.9 | <.001 |
| R dPMC | 27 | -7 | 65 | 6.4 | <.001 |
| L dPMC | -25 | -8 | 56 | 3.7 | <.001 |
| R MFG | 35 | 44 | 15 | 3.4 | .046 |
| L MFG (BA 46) | -28 | 33 | 30 | 3.9 | .028 |

The HMAX C1 model confirmed that the low-level features were successfully represented in the analysis, acting as a baseline (Fig. 4b). The results revealed strong representational similarity effects predominantly in early visual cortex and adjacent occipito-temporal regions. Bilateral Calcarine cortex (R: *t*(24) = 10.4; L: *t*(24) = 10.7, *p* < .001) and Lingual Gyrus (R: *t*(24) = 8.3; L: *t*(24) = 8.5, *p* < .001) showed robust effects. Significant effects were also observed in bilateral Superior Occipital Gyrus (SOG; R: *t*(24) = 15.7; L: *t*(24) = 13.2, *p* < .001) and bilateral Middle Occipital Gyrus (MOG; R: *t*(24) = 8.7; L: *t*(24) = 11.0, *p* < .001).

Beyond early visual regions, the right Fusiform Gyrus also showed significant effects (R: *t*(24) = 7.7 *p* < .001), along with the Right Cuneus (*t*(24) = 11.6, *p* < .001). In addition, direction-related similarity explained by low-level features was observed in right MT/V5 (*t*(24) = 4.8, *p* < .001).

Overall, the HMAX C1 model revealed widespread effects across occipital and ventral temporal regions, indicating that low-level visual structure contributes significantly to representational similarity patterns in early and mid-level visual areas. The peak coordinates, *t*-values, and corrected *p*-values thresholded with the TFCE method (*p* <.05, *z* = 1.65) for all clusters are reported in Table 4.

**Table 4.** The region labels, peak MNI coordinates, *t*- and *p*-values of multiple regression RSA results of the HMAX C1 model. R: right, L: left, SOG: superior occipital cortex, MOG: middle occipital cortex, MT: middle temporal area.

|  | MNI Coordinates |  |  |  |  |
| --- | --- | --- | --- | --- | --- |
| Region (AAL3) | x | y | z | $t(24)$ | $p$ |
| R Calcarine | 21 | -94 | 0 | 10.4 | <.001 |
| L Calcarine | -6 | -96 | 1 | 10.7 | <.001 |
| R Lingual | 3 | -82 | -6 | 8.3 | <.001 |
| L Lingual | -8 | -85 | -12 | 8.5 | <.001 |
| R SOG | 23 | -97 | 14 | 15.7 | <.001 |
| L SOG | -8 | -102 | 12 | 13.2 | <.001 |
| R MOG | 41 | -83 | 5 | 8.7 | <.001 |
| L MOG | -13 | -102 | 9 | 11 | <.001 |
| R Fusiform | 29 | -76 | -12 | 7.7 | <.001 |
| R Cuneus | 18 | -98 | 12 | 11.6 | <.001 |
| R MT | 45 | -71 | 0 | 4.8 | <.001 |

The results of the action type model also yielded widespread representation in AON, including occipital, occipito-temporal, temporal, somatomotor, parietal, and frontal areas, with the strongest representation in the occipital and occipito-temporal cortices (see Supplementary File 1- Fig S2). Furthermore, for the motor model, bilateral Precentral (M1; R; *t*(24) = 4.7; L; *t*(24) = 2.0, *p* <.001), and Postcentral (S1; R; *t*(24) = 4.6, *p* <.001 L; *t*(24) = 3.1 , *p* = .002), as well as right Inferior Parietal Sulcus (IPS; R; *t*(24) = 4.8, *p* <.001) areas showed significant representations (see Supplementary File 1- Figure S3). Finally, the tool model and the random model did not yield any significant clusters in the TFCE thresholded map (p > .05). The unthresholded one sample t-test results of the tool model can be found in Supplementary File 1- Figure S4.

## Discussion

Although everyday actions are rich in direction information, previous studies have not investigated direction selectivity with naturalistic actions, but have used PLDs and simple moving dots or gratings. The present study addressed this gap by investigating the neural basis of action direction selectivity using naturalistic, repetitive, bidirectional action videos and multivariate fMRI analyses. First, we employed MVPA to identify brain regions that contained sufficient information to discriminate between action directions (i.e., decodability; Peelen & Downing, 2023; Popal et al. 2020). While informative, decoding analysis may conflate direction with correlated features and/or underrepresent certain direction-selective areas (Diedrichsen & Kriegeskorte, 2017; Popal et al. 2020). To address this issue, we then applied multiple regression RSA, which enables testing the unique contribution of direction information after accounting for low-level, motor, and action-related effects (Freund et al. 2021; Popal et al. 2020; Xue et al. 2013). Because RSA directly compares neural similarity matrices with model-derived structures, it offers greater sensitivity and localizes representationally relevant regions more precisely than MVPA (Freund et al. 2021; Weaverdyck et al. 2020). In this framework, MVPA provided a critical baseline by establishing the decodability of action directions, while RSA refined these results by isolating direction-specific representations. Together, these complementary methods allowed us to robustly characterize the neural coding of action direction.

Results have revealed strong decoding and representation of action direction within the action observation network (AON), including occipital, occipito-temporal, parietal, motor, and frontal areas bilaterally. These findings extend prior work on motion perception and biological motion processing, providing novel evidence that action direction is represented not only in low-level sensory regions (i.e., visual cortex and MT), but also in higher-level action-related areas when ecologically valid, naturalistic action stimuli are used.

Consistent with previous studies in macaques and humans, strong direction decoding and representation in early visual regions (i.e., calcarine cortex, lingual area, MOG, SOG, MT/V5) were observed (Albright, 1984; Hong et al. 2012; Stoppel et al. 2011; Vaina et al. 2001; Vangeneugden et al. 2014; Wurm & Erigüç, 2025; Ziccarelli et al. 2024). Additionally, the multiple regression RSA confirmed that these effects could not be explained solely by low-level visual features captured by the HMAX C1 model, suggesting direction-specific neural coding. The involvement of left LOC further supports that occipito-temporal areas contribute not only to object and form processing (Eger et al. 2007; Snow et al. 2015) but also to the integration of motion cues relevant for action interpretation (Vaina et al. 2001; Vangeneugden et al. 2014; Weiner & Grill-Spector, 2011). Together, these results suggest that occipital and occipito-temporal cortices are essential for encoding direction across a range of visual and action contexts.

Results have revealed direction-selective representations within the parietal cortex, specifically in both the inferior parietal sulcus (IPS) and the superior parietal lobule (SPL). The IPS is a key multimodal area that integrates visual information with spatial properties, and it has been strongly implicated in action observation and imitation (Caspers et al. 2010; Fogassi & Luppino, 2005). This integrative role makes it a plausible candidate for representing action direction, as accurate interpretation of an observed action often requires linking visual motion cues with their spatial orientation and trajectory (Karnath, 1997). Within the SPL, our findings are consistent with previous work showing that the superior parieto-occipital cortex (SPOC) decodes the movement direction of point-light display (PLD) hand movements using MVPA (Ziccarelli et al. 2024). More broadly, the involvement of intraparietal areas may indicate abstract representations of direction, in line with evidence from both macaque and human studies (Freedman & Assad, 2006; Ziccarelli et al. 2024). Supporting this view, electrophysiological research has shown that the posterior parietal cortex (PPC), which includes the lateral intraparietal area (LIP) and the parietal reach region (PRR), contains neurons tuned to motion direction (Andersen & Buneo, 2002).

In addition to parietal regions, results have revealed that the sensorimotor regions robustly represented action direction: SMA, dorsal premotor, primary motor, and primary somatosensory areas. The results of the dorsal premotor cortex are consistent with a previous study that found that the direction of an abstract animation (i.e., right-to-left and left-to-right) was decoded in the right dorsal premotor area and MT/V5 (Wurm & Erigüç, 2025). Moreover, a study on hand movement direction during motor planning and execution reported that SMA and parietal areas are engaged in planning, whereas motor and premotor cortices decode movement direction during execution (Combrisson et al. 2024). Similarly, in the reaching task (i.e., left or right), direction information was represented within superior parietal and premotor areas (Caceres et al. 2024). Another study involved the wrist and forearm movements of 8 directions and found that the primary motor cortex, supplementary motor area, and premotor cortex were selective to movement directions of the hand (Cowper-Smith et al. 2010). Furthermore, a similar study with wrist movements of monkeys has found that the neurons in the ventral premotor area were directionally tuned independent of the position of the posture of the hand (Kakei et al. 2001). Although these works addressed action execution rather than observation, our findings converge, as action observation and execution are closely linked within overlapping neural circuits, especially in parietal and motor cortices (Culham & Valyegar, 2006; Grezes & Decety, 2001; Hardwick et al., 2018; Vignemont & Haggard, 2008). Linking this relationship with execution and observation, it has been frequently reported in the literature that observing touch and hand related actions recruits somatosensory cortex (Avikainen et al., 2002; Blakemore et al., 2005; Gazzola & Keysers, 2008; Keysers et al., 2004; Meyer et al., 2011; Rossi et al., 2002; Schaefer et al., 2009), as well as primary motor cortex (Dushanova & Donoghue, 2010; Hari et al., 1998; Rassi et al., 2024). In fact, Tkach et al. (2007) demonstrated that neurons in M1 maintain nearly identical preferred directions during both the execution and passive observation of familiar tasks. M1 neurons are capable of representing the specific trajectory and direction of observed actions in a way that is consistent with the motor system’s own structure (Dushanova & Donoghue, 2010; Tkach et al., 2007). Direction preference of the neurons are not limited to the motor cortex, but somatosensory cortex as well, although this was initially measured only for tactile stimuli (Costanzo & Gardner, 1980). Furthermore, it has been reported that neurons in S1 represent the direction of tactile motion in a shape-invariant manner, maintaining a consistent preferred direction across different stimulus types (Pei et al., 2010). Crucially, the sensitivity of these neurons matched human psychophysical performance in direction discrimination tasks. These findings add physiological support to the current study that parietal and somatomotor regions can robustly represent the direction of observed actions, potentially by recruiting these high-level motion processing mechanisms.

Although previous work has revealed direction selectivity at multiple levels of the action observation network (AON) from low-level visual regions such as V1-V4 and MT/V5 (Albright, 1984; Hong et al. 2012; Stoppel et al. 2011) to higher-order parietal and premotor cortices (Freedman & Assad, 2006; Wurm & Erigüç, 2025; Caceres et al. 2024), these studies have typically highlighted specific regions rather than demonstrating a broad, network-wide representation of direction information. The present study revealed that direction information of complex, bidirectional actions is represented across multiple levels of the action observation network (AON), reflecting the multi-layered nature of perceptual, motor, and cognitive processes involved in interpreting complex actions. Such widespread engagement may be necessary because actions encountered in daily life are rarely simple or unidirectional but instead involve rich spatial dynamics that must be integrated for accurate perception and understanding. Importantly, this study shows that direction selectivity is not restricted to early visual regions, as in the previous visual direction selectivity studies, but higher-order parietal and frontal regions also contribute when the observed actions are more naturalistic and complex, highlighting the multilayered coding of direction within the human brain. These results might suggest that direction information may be represented hierarchically, in early visual regions such as V1/V2 and MT in a sensory manner, and extended into more abstract representations in higher-order areas such as the premotor and PMv, where the direction of actions may be categorically interpreted in terms of their meaning or goal.

Several limitations of the current study should be acknowledged. One limitation concerns the stable positioning of the on-screen response texts. Because participants indicated action direction using their thumbs and response options appeared in fixed spatial locations, the task may have induced motor preparation or imagery during action observation. Accordingly, the strong decoding accuracy in primary motor and somatosensory cortices during MVPA may partly reflect motor planning related to the upcoming button press rather than purely perceptual representations of direction. In the present study, we addressed this concern by conducting a multiple regression RSA that explicitly modeled potential motor confounds. Nevertheless, future studies could further reduce motor-related influences by randomizing response locations across trials or by systematically controlling the responding fingers or hands to minimize consistent motor preparation effects.

Additionally, the middle button was not pressed with the same thumb across all participants, introducing variability in motor responses that, although accounted for by subject-specific motor models in the RSA, may have added residual motor-related noise. Another limitation is that our stimuli were limited to a set of three bidirectional movement dimensions. Using naturalistic actions that are non-repetitive, either bidirectional or unidirectional, could provide valuable insight into whether the observed representational patterns generalize beyond repetitive trajectories. Additionally, because our stimuli consisted of only right-hand actions and all analyzed participants were right-handed, future studies could systematically manipulate both the handedness of the observed actions and the participants to determine whether handedness modulates the neural representation of action direction. — Furthermore, using communicative action stimuli such as waving a hand or handshaking could reveal the processing of directions regarding socially meaningful stimuli. A promising approach would also be to employ more naturalistic, real-world setups combined with EEG recordings, where depth and three-dimensional motion cues can be captured with higher ecological validity and temporal precision. Despite these limitations, the present study offers important insights into the global processing of direction information in the action observation network for complex, repetitive, and naturalistic actions.

## Conclusion

The present study provides novel evidence that direction selectivity during naturalistic action observation is not limited to early visual cortices but is broadly distributed across the action observation network. Using complementary multivariate approaches, MVPA and RSA, we demonstrated that direction information is represented in occipital, temporal, parietal, and frontal regions. Importantly, these results extend prior work based on simplified or point-light stimuli by showing that complex, repetitive, and ecologically valid actions demonstrate widespread, global representations of direction.

Taken together, these findings highlight the distributed and integrative nature of direction coding in the human brain, supporting the view that action direction is a fundamental property represented at multiple levels of the action observation network. This broad representation may allow the brain to flexibly interpret action trajectories in naturalistic contexts, bridging low-level motion processing with higher-order action understanding and prediction.

## Acknowledgements

The authors would like to thank Ece Altınbaş for her contributions in stimulus conceptualization and creation, and Cenk Günsel for his contributions in stimulus recording. Additionally, the authors appreciate the undergraduate research assistants who helped with fMRI data collection. The authors would also like to thank the Turkish Health Institutes Presidency (TUSEB) for funding this study (Project No: 37734). The first author was supported by TUBITAK-BIDEB 2210-National Graduate Scholarship Programme. The preprocessing of the fMRI data was fully performed at TUBITAK ULAKBIM, High Performance and Grid Computing Center (TRUBA resources).

## Author Contributions

**Zelal Eltaş**: Writing – original draft, Investigation, Visualization, Software, Methodology, Formal analysis, Conceptualization, Funding acquisition. **Murat B. Tunca**: Writing – review & editing, Formal analysis, Supervision, Methodology, Conceptualization, Funding acquisition. **Burcu A. Urgen**: Writing – review & editing, Supervision, Methodology, Conceptualization.

## Data and Code Availability

Data will be made available upon request. Stimulus set and analysis codes of the current study are available at the Open Science Framework’s website at https://osf.io/9sqyd/

## Notes

### Competing Interest Statement

The authors have declared no competing interest.

### Summary of Updates

In the introduction, the aim of the current study was clarified. In the results section, the preprocessing steps were provided in greater detail. Additionally, the RSA results changed because the underlying models were updated; specifically, a "Tool" model was added, and the "Motor Response" model was divided into two sub-models based on which hand the participants used. Following the inclusion of these models, the Results and Methods sections were revised, and Figures 2 and 4 were updated accordingly. Finally, in the discussion, the involvement of the somatomotor cortex was discussed in more detail, and the limitations section was expanded.

