## Supplementary File 1 for "Direction Selectivity in Naturalistic Action Observation: Distributed Representations Across the Action Observation Network"

**Title:** Direction Selectivity in Naturalistic Action Observation: Distributed Representations Across the Action Observation Network

**Authors:** Zelal Eltaş, Murat B. Tunca, and Burcu A. Urgen

**Fig. S1** Method to define which thumb participants used for the middle button press. Example results from two participants, one of whom used their right thumb (a) for the middle button press and the other (b) used their left thumb.

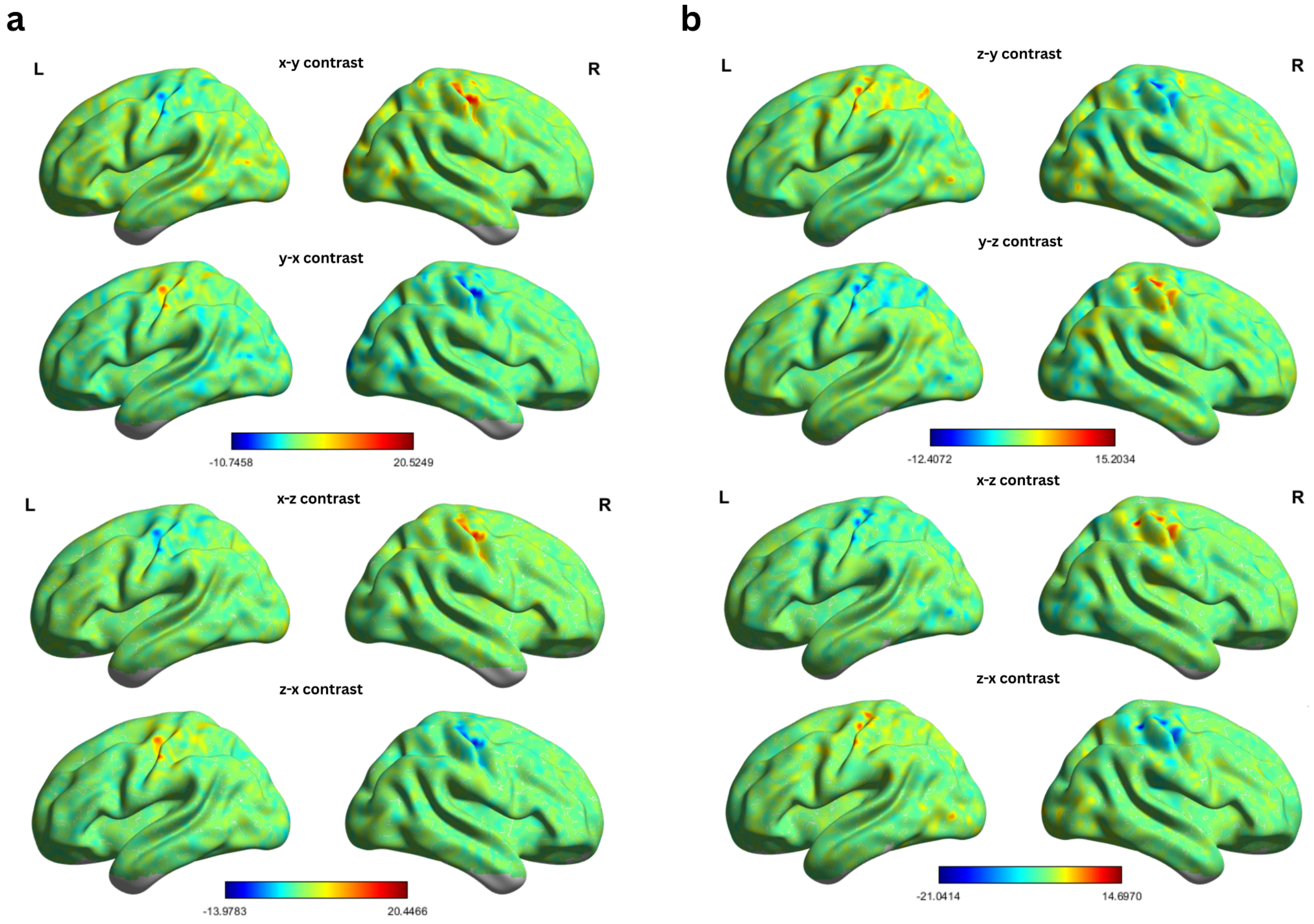


It is known that participants used their right thumb for front-back actions, and their left thumb for right-left actions. To determine which thumb was planned to use for the middle button press, the GLM data used for the MVPA was examined. The GLM consists of x, y, and z actions, which correspond toright-left, up-down, and front-back actions, respectively. Pairs of x–y, y–x, x–z, z–x, y–z, z–y were determined for GLM contrast analysis. Across all subjects, contralateral primary motor and somatosensory activations were observed. For example, because x actions were performed with the left thumb and z actions with the right thumb, the x–z contrast showed activation in right M1 and S1, while the reverse contrast showed the opposite pattern. Importantly, these results originate from the observation phase of the study, not the response period.

To illustrate the inference method used about the thumb participants plan to use, two representative participants were demonstrated in Figure S1. In panel a, the x–y contrast (left button minus middle button) shows activation in the right precentral area, with corresponding negative t-values in the left hemisphere. This pattern closely resembles the x–z contrast (left button minus right button), suggesting that the participant used their right hand to press the middle button. In contrast, Panel b shows an opposite pattern: the z–y contrast (right button minus middle button) reveals left precentral activation, with negative trends in the corresponding contralateral region. This pattern is opposite to that observed in the x–z and z–x contrasts, concluding that the participant used their left hand when planning to press the middle button.

**Fig. S2** One-sample t-test multiple regression RSA results, thresholded with TFCE-based Monte Carlo simulation with 10,000 permutations ($p < 0.05$, $z = 1.65$) for the action model, when accounting for HMAX C1, direction, motor, tool, and random models. Colorbar shows t-values.

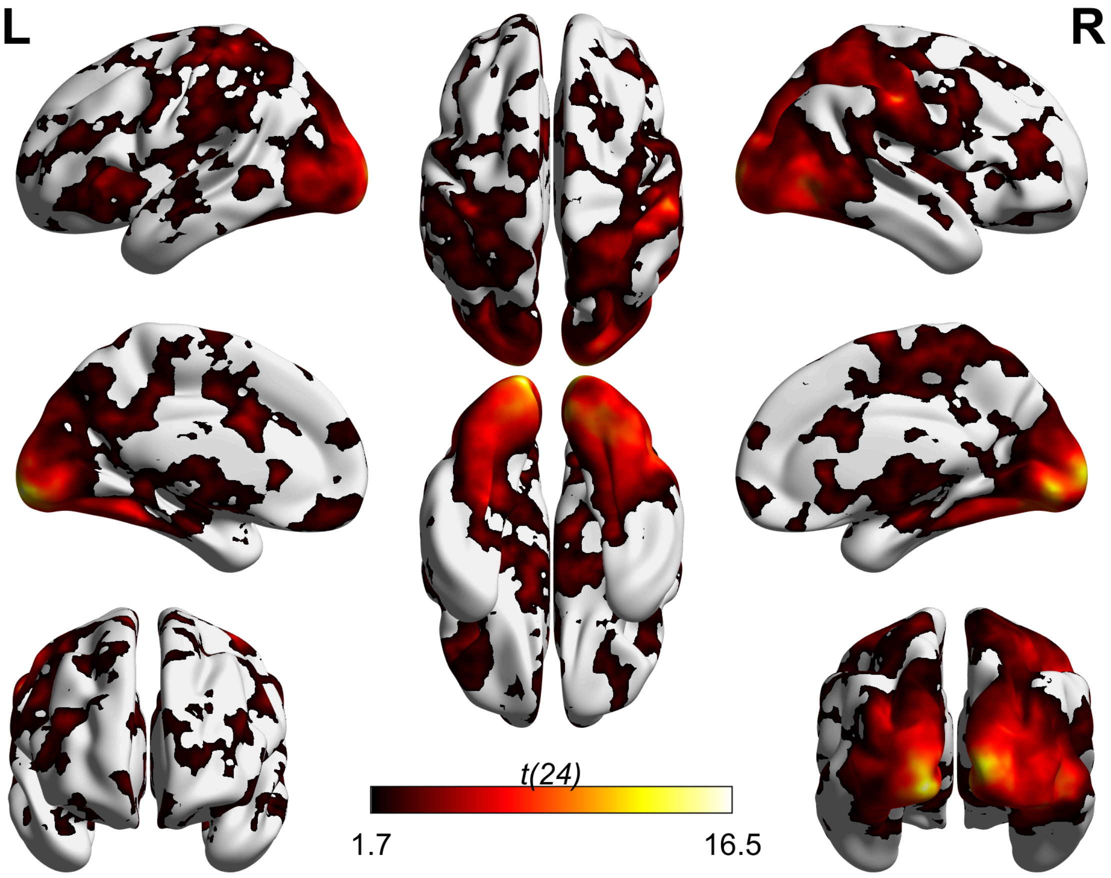

**Fig. S3** One-sample t-test multiple regression RSA results, thresholded with TFCE-based Monte Carlo simulation with 10,000 permutations ($p < 0.05$, $z = 1.65$) for the motor model, when accounting for HMAX C1, direction, action, tool, and random models. Colorbar shows t-values.

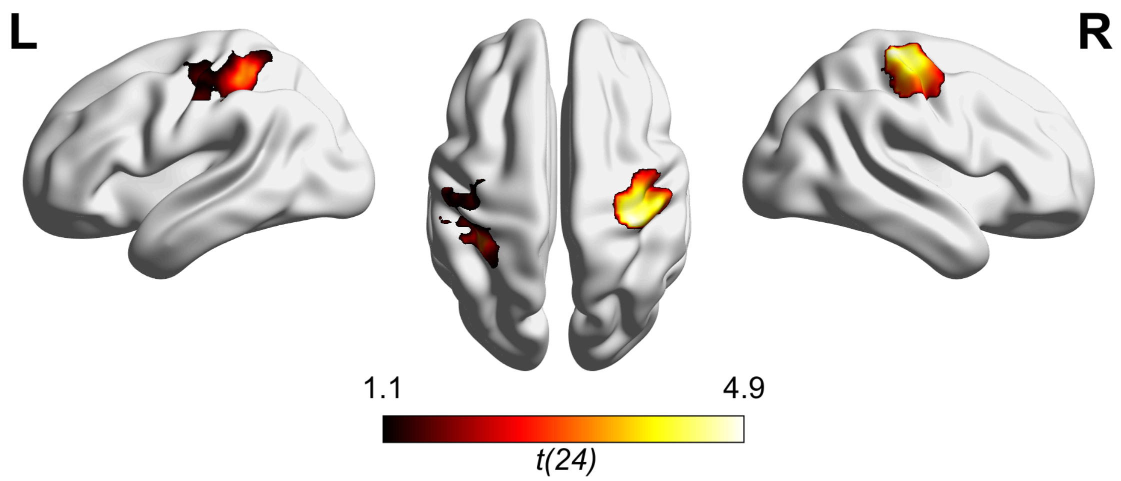

**Fig S4** Unthresholded one-sample t-test results, for the tool model, when accounting for HMAX C1, direction, motor, action, and random models. Colorbar shows t-values.

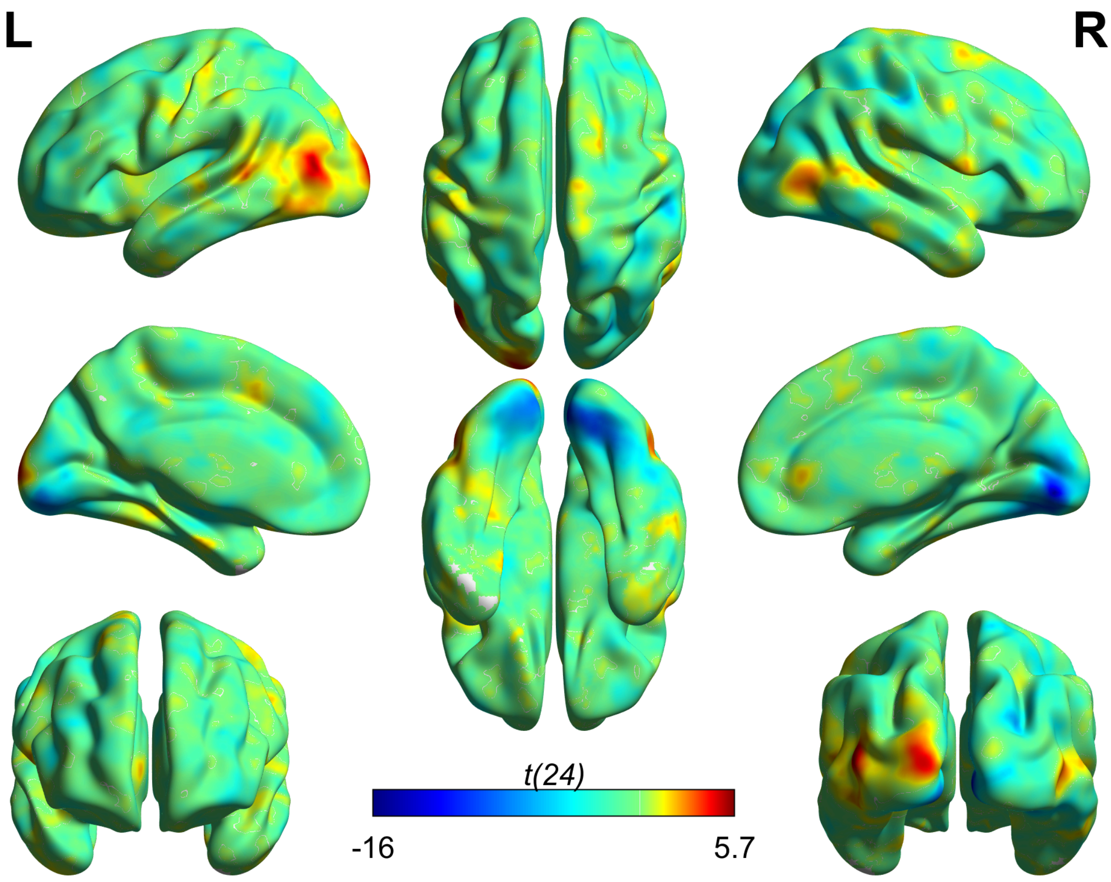


The unthresholded group t-test map shows tool-sensitive areas such as the posterior Middle Temporal Gyrus and Inferior Temporal Cortex, with a dominance in the left hemisphere since the participants were right-handed (Lewis, 2006; Mruczek et al., 2013; Orban & Caruana, 2014; Ramayya et al., 2009).
