## Supplementary figures and images for "Direction Selectivity in Naturalistic Action Observation: Distributed Representations Across the Action Observation Network"

### Supplementary File 2

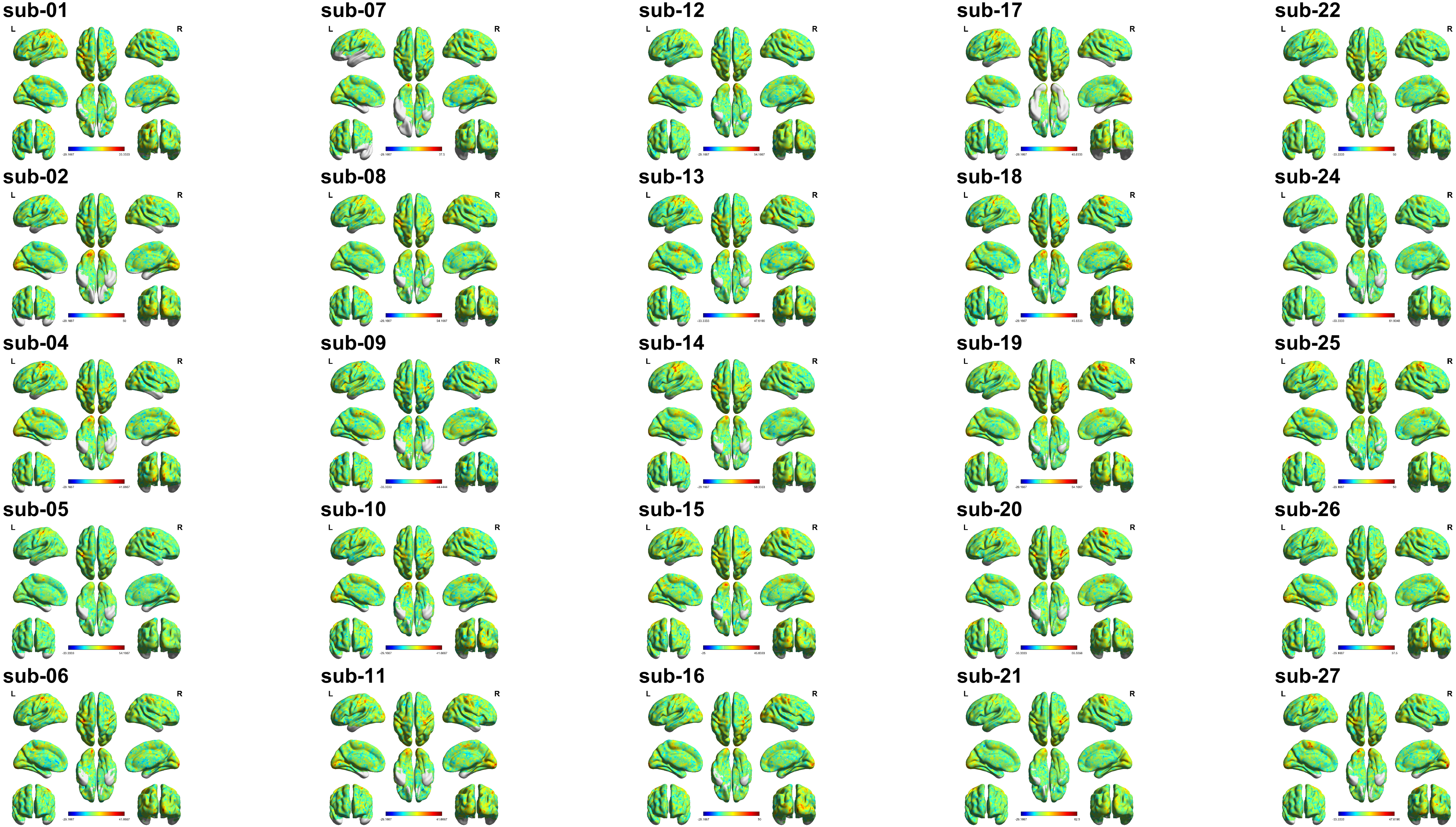
